# Golgi outposts locally regulate microtubule orientation in neurons but are not required for the overall polarity of the dendritic cytoskeleton

**DOI:** 10.1101/866574

**Authors:** Sihui Z. Yang, Jill Wildonger

## Abstract

Microtubule-organizing centers (MTOCs) often play a central role in organizing the cellular microtubule networks that underlie cell function. In neurons, microtubules in axons and dendrites have distinct polarities. Dendrite-specific Golgi outposts, in particular multi-compartment outposts, have emerged as regulators of acentrosomal microtubule growth, raising the question of whether outposts contribute to establishing the overall polarity of the dendritic microtubule cytoskeleton. The *cis*-Golgi matrix protein GM130 has roles in both the MTOC activity of Golgi and in connecting Golgi compartments to form multi-compartment units. Using a combination of genetic approaches and live imaging in a Drosophila model, we found that GM130 is not essential for the overall polarity of the dendritic microtubule cytoskeleton. However, the mislocalization of multi-compartment Golgi outposts to axons disrupts the uniform orientation of axonal microtubules. This suggests that outposts have the capacity to influence microtubule polarity and, as our data indicate, likely do so independently of microtubule nucleation mediated by the γ-tubulin ring complex (γ-TuRC). Altogether, our results are consistent with the model that multi-compartment Golgi outposts may locally influence microtubule polarity, but that outposts are not necessary for the overall polarity of the dendritic microtubule cytoskeleton.

## INTRODUCTION

Proper neuronal structure and function depends on the underlying microtubule cytoskeleton, which is uniquely organized in axons and dendrites. The compartment-specific orientation of microtubules is thought to contribute to the specific morphologies and functions of these compartments. In axons, microtubules are uniformly oriented with their plus-ends positioned distal to the cell body. In contrast, microtubule polarity in dendrites is mixed to varying degrees, and the percentage of microtubules of a particular orientation differs locally within the dendritic arbor. How the distinctly organized cytoskeletons in axons and dendrites are created and maintained remains an open question. Microtubules are often generated at and organized by cellular structures called microtubule-organizing centers (MTOCs) that anchor and stabilize microtubules and support microtubule nucleation (Sanchez and Feldman, 2017; Wu and Akhmanova, 2017). The centrosome is one example of a well-studied MTOC. Although neurons have a centrosome, recent work indicates that the neuronal centrosome does not have a major role in either generating or anchoring dendritic or axonal microtubules (Nguyen et al., 2011; Sanchez-Huertas et al., 2016; Stiess et al., 2010). The centrosome, however, is not the only organelle that functions as an MTOC (Sanchez and Feldman, 2017; Wu and Akhmanova, 2017). The Golgi apparatus and non-conventional Golgi structures such as Golgi elements and Golgi outposts have emerged as potential MTOCs in several cell types, including epithelia, muscles, and neurons (Martin and Akhmanova, 2018; Rios, 2014; Sanders and Kaverina, 2015). Unlike centrosomes, which support the nucleation of microtubules radially, Golgi-based microtubule nucleation can create asymmetric microtubule arrays (Efimov et al., 2007; Zhu and Kaverina, 2013). Such asymmetric Golgi-derived microtubule arrays have also been shown to contribute to cell polarity (Rios, 2014; Zhu and Kaverina, 2013). Thus, Golgi in neurons have the potential to shape the polarity of the microtubule cytoskeleton, and influence neuronal polarity, by selectively seeding or stabilizing microtubules in a particular orientation.

In developing flies and mammals, many neurons have dendrite-specific Golgi mini-stacks called "outposts" that are distinct from the somatic Golgi (Gardiol et al., 1999; Horton and Ehlers, 2003; Liu et al., 2017; Pierce et al., 2001; Rao et al., 2018; Tann and Moore, 2019; Ye et al., 2007). Golgi and Golgi outposts are excluded from axons. The term Golgi outposts is used to refer generally to a heterogenous population of dendritic Golgi composed of one or more compartments. In studies using fluorescently tagged End-binding 1 (EB1::GFP), which marks growing microtubule ends, Golgi outposts have been shown to correlate with microtubule growth initiation sites in developing dendrites (Ori-McKenney et al., 2012; Yalgin et al., 2015; Zhou et al., 2014). This correlation has led to the model that Golgi outposts function as MTOCs that support acentrosomal nucleation and the directional growth of microtubules during dendrite branch formation (Delandre et al., 2016). Decreasing the levels of proteins involved in microtubule nucleation, such as γ-tubulin and its interactors centrosomin (cnn, the fly ortholog of CDK5RAP2) and pericentrin-like protein (plp, the Drosophila ortholog of AKAP450), disrupts the correlation between microtubule growth initiation sites and outposts and perturbs dendrite branch growth (Ori-McKenney et al., 2012; Yalgin et al., 2015). The link between microtubule growth and Golgi outposts relies on the compartmental organization of outposts: multi-compartment dendritic Golgi outposts are more likely than single-compartment outposts to correlate with microtubule growth start sites (Zhou et al., 2014) and dragging individual Golgi compartments into the axon is not sufficient to alter axonal microtubule organization (Nguyen et al., 2014). The dendrite-specific localization of Golgi outposts and their MTOC potential raises the possibility that Golgi outposts, and in particular multi-compartment outposts, may play a role in establishing and/or maintaining the unique polarity of the dendritic microtubule cytoskeleton; this idea, however, has not been tested.

Here we investigated whether Golgi outposts are necessary for the compartment-specific orientation of dendritic microtubules. In particular, we asked whether outposts are necessary for the formation of the minus-end-distal microtubules that are specific to the dendritic cytoskeleton. To test this notion, we leveraged a combination of genetics and live imaging and used the class IV dendritic arborization (da) neurons in Drosophila as a model. In the class IV da neurons, dendritic microtubules are predominately oriented with their minus-ends distal to the cell body, providing an advantageous paradigm to detect changes in microtubule orientation. If Golgi outposts regulate the overall polarity of the dendritic microtubule cytoskeleton, then blocking the MTOC activity of Golgi should disrupt the stereotyped minus-ends-distal orientation of dendritic microtubules. We targeted the *cis*-Golgi matrix protein GM130, which has two key roles: first, work done in mammalian cells has shown that GM130 recruits AKAP450, which in turn recruits protein complexes that nucleate, tether, and stabilize microtubules (Hurtado et al., 2011; Rivero et al., 2009; Roubin et al., 2013; Wu and Akhmanova, 2017). Second, GM130 is needed for proper Golgi structure and connects Golgi compartments to form multi-compartment units, including outposts (Barr et al., 1997; Kondylis et al., 2005; Liu et al., 2017; Lowe, 2019; Nakamura et al., 1995; Zhou et al., 2014). Strikingly, we found that the global orientation of dendritic microtubules is unaffected by the loss of GM130 or the fly AKAP450 ortholog plp. This suggests that Golgi outposts do not have an essential role in establishing the overall polarity of the dendritic microtubule cytoskeleton. Nonetheless, it is still possible that outposts have a role in locally or dynamically regulating the microtubule cytoskeleton. In support of this idea, we found that Golgi outposts that mislocalized to axons altered microtubule polarity. The effect of the ectopic outposts on microtubule polarity was independent of γ-TuRC and its regulator cnn but dependent on the formation of multi-compartment Golgi outposts, which could be induced by increasing GM130 levels. Based on our data and the work of others, we propose that multi-compartment Golgi outposts may act to locally remodel the microtubule cytoskeleton during dendrite arborization, but that Golgi outposts do not play a significant role in creating or maintaining the unique overall polarity of the dendritic microtubule cytoskeleton.

## RESULTS

### Removing GM130 reveals Golgi outposts are not essential to the overall polarity of the dendritic microtubule cytoskeleton

To determine whether Golgi outposts play a critical role in determining the unique polarity of the dendritic microtubule cytoskeleton, we turned to the class IV da neurons in Drosophila as a model. The da neurons are an ideal model as they are easily accessible for live imaging, the orientation of microtubules in the dendritic microtubule cytoskeleton is well-defined, and there is a wealth of tools for labeling and manipulating Golgi, microtubules, and microtubule regulators. In the class IV da neurons, Golgi compartments are present in the cell body and throughout the dendritic arbor, but most are found close to the cell body (Figure 1, A-C). Indeed, GalNacT2-positive *trans* Golgi compartments are seldom present beyond ~100 µm of the cell body. Next, we quantified the number of multi-compartment outposts in the arbor. Consistent with previous work, we defined multi-compartment Golgi outposts as units with at least two compartments (Zhou et al., 2014). Here, we used markers of the *medial* and *trans* Golgi, namely ManII::GFP and GalNacT2::TagRFP, respectively. Fewer than half the ManII-positive Golgi outposts colocalized with GalNacT2, indicating ManII::GFP predominantly labels single-compartment Golgi outposts (Figure 1D). In contrast, the majority of GalNacT2-positive outposts colocalized with ManII::GFP (Figure 1D), which suggests that most multi-compartment Golgi outposts cluster relatively close to the cell body.

**Fig. 1.**
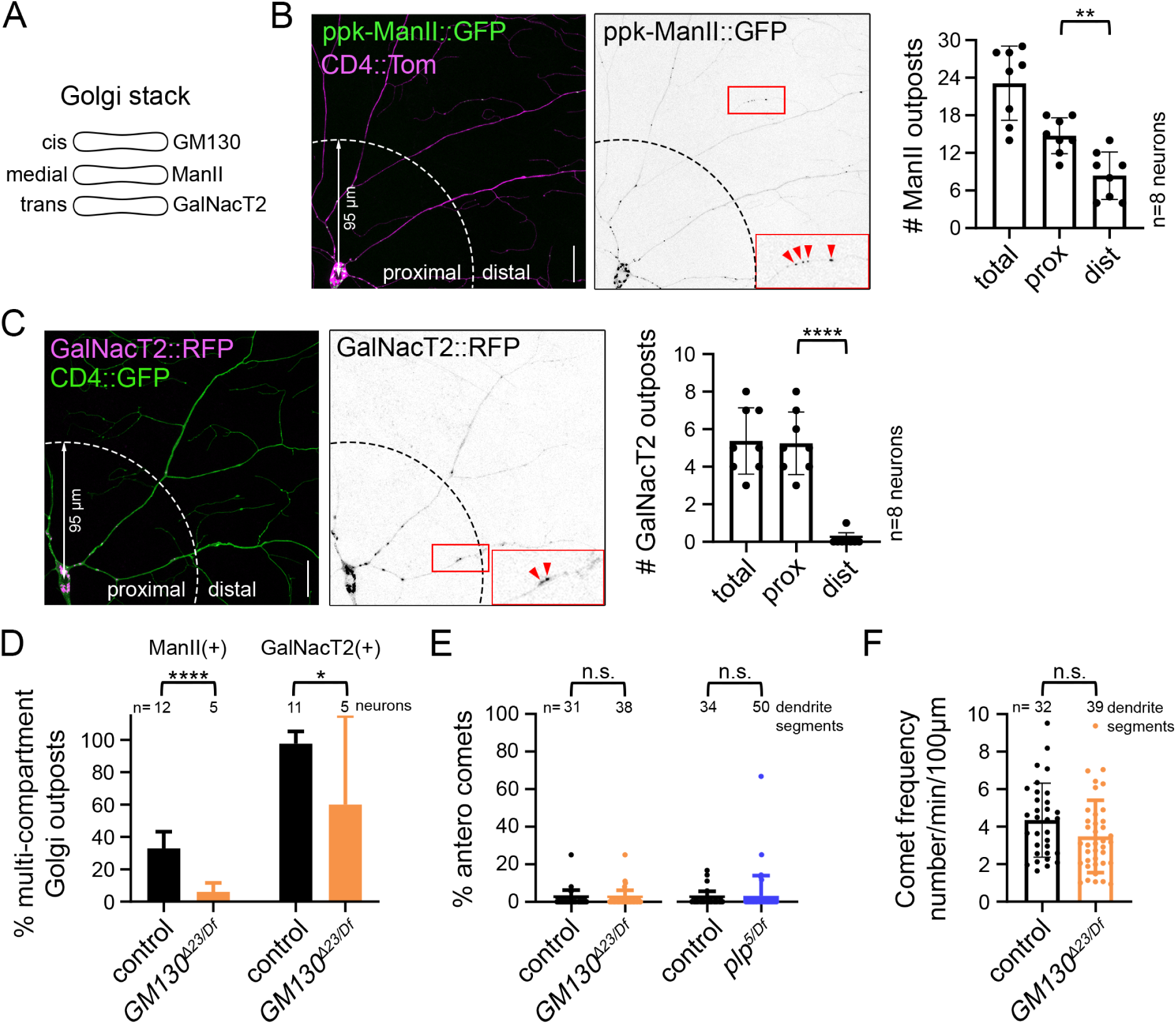
Global dendritic microtubule polarity does not rely on the *cis*-Golgi protein GM130. (A) Cartoon showing the compartmental distribution of GM130, ManII and GalNacT2 in a Golgi stack. (B-D) ManII::GFP-positive outposts (B, left graph in D) are present throughout the class IV da neuron dendritic arbor, but fewer than half of these outposts are multi-compartment units. Multi-compartment outposts are defined as those that have overlapping ManII::GFP and GalNacT2::TagRFP signal. In contrast to ManII::GFP-positive outposts, GalNacT2::TagRFP-positive outposts (C, right graph in D) cluster in the proximal arbor and nearly all GalNacT2::TagRFP-positive outposts are multi-compartment. Eliminating GM130 reduces the percentage of outposts that are multi-compartment in the proximal arbor. The proximal arbor (prox) encompasses a radius of 95 µm from the cell body; distal (dist) is beyond this radius. The left side of the graph represents the fraction of ManII::GFP-positive outposts that are multi-compartment and the right side of the graph represents GalNacT2::TagRFP-positive outposts that are multi-compartment. \**P*=0.05–0.01, \*\**P*=0.01–0.001, and ****P<0.0001; Student’s unpaired t tests. (E) Dendritic microtubule polarity is not affected by the loss of either GM130 (left) or plp (right). Antero = anterograde. n.s.=not significant, Mann-Whitney test. (F) EB1::GFP comet frequency is normal in the dendrites of neurons lacking GM130. n.s.=not significant, Student’s unpaired t-test. Microtubule polarity and EB1::GFP comet frequency were quantified in the proximal dendritic arbor which contained the majority of multi-compartment outposts. Scale bars: 25 µm. All data are mean ± standard deviation.

To determine whether Golgi outposts function to create or maintain the dendrite-specific orientation of microtubules, we analyzed microtubule polarity in neurons in which we eliminated the *cis*-Golgi matrix protein GM130. GM130 has the potential to both recruit the protein machinery for MTOC activities (microtubule nucleation, anchoring, and stabilization) and contribute to forming multi-compartment Golgi units (Barr et al., 1997; Kondylis et al., 2005; Liu et al., 2017; Lowe, 2019; Martin and Akhmanova, 2018; Nakamura et al., 1995; Sanders and Kaverina, 2015; Zhou et al., 2014). We found that the percentage of multi-compartment outposts decreases when GM130 is absent, which supports the model that GM130 participates in connecting Golgi compartments in neurons (Figure 1D)(Zhou et al., 2014). To read-out microtubule orientation, we used EB1::GFP, whose binding to growing microtubule ends produces a comet-like trajectory. The majority of microtubule growth occurs at plus-ends, which also grow faster than minus-ends, enabling a clear distinction of plus-and minus-end growth and thus microtubule polarity (Feng et al., 2019). As previously reported, in control neurons dendritic microtubules are oriented predominantly minus-ends-distal (Figure 1E). Strikingly, eliminating GM130 had no effect on the overall polarity of microtubules within the proximal dendritic arbor where multi-compartment Golgi outposts clustered (Figure 1E). The overall frequency of microtubule growth was also unaffected by the loss of GM130 (Figure 1F). In mammalian cells, GM130 affects microtubule organization through the recruitment of AKAP450, whose fly ortholog is plp (Martin and Akhmanova, 2018; Martinez-Campos et al., 2004; Rios, 2014; Sanders and Kaverina, 2015). Plp in fly neurons has likewise been implicated in the MTOC activity of Golgi outposts (Ori-McKenney et al., 2012). Similar to the loss of GM130, eliminating plp had no effect on dendritic microtubule polarity (Figure 1E). Thus, the results of our experiments indicate that Golgi outposts do not play an essential role in creating or maintaining the overall polarity of the dendritic microtubule cytoskeleton.

### Loss of GM130 significantly reduces misoriented microtubules in *nudE*^−^ axons

The results of our GM130 loss-of-function experiments in da neurons suggest that the overall polarity of the dendritic microtubule cytoskeleton does not depend on Golgi outposts. Microtubule polarity, however, varies within the dendritic arbors of da neurons (Stone et al., 2008), and it is possible that Golgi outposts are nevertheless sufficient to locally influence microtubule polarity as previously reported (Delandre et al., 2016; Ori-McKenney et al., 2012; Yalgin et al., 2015). To test the idea that outposts have the capacity to affect microtubule polarity, we used a paradigm in which Golgi outposts are ectopically localized to axons, a compartment from which outposts are normally excluded and in which microtubules are uniformly oriented with their plus-ends distal. If Golgi outposts are capable of regulating microtubule organization, we reasoned that their ectopic presence in axons should disrupt the uniform plus-ends-distal array of axonal microtubules. To ectopically localize Golgi outposts to axons, we relied on mutations that disrupt the activity of the molecular motors dynein and kinesin-1, which transport outposts (Arthur et al., 2015; Kelliher et al., 2019; Lin et al., 2015; Ye et al., 2007; Zheng et al., 2008). While disrupting the activity of either motor results in Golgi outposts invading axons, we and others have previously shown that these ectopic outposts do not always correlate with a change in axonal microtubule polarity (Kelliher et al., 2019; Nguyen et al., 2014; Ye et al., 2007). Analyzing these different motor mutants and the outposts in their axons enables us to determine whether Golgi outposts are sufficient to affect microtubule polarity and to then identify the factors that are essential for this activity.

The loss of dynein activity alters both Golgi outpost localization and axonal microtubule polarity (Arthur et al., 2015; del Castillo et al., 2015; Klinman et al., 2017; Rao et al., 2017; Zheng et al., 2008). We first asked whether the misoriented microtubules in dynein loss-of-function neurons depend on the ectopic Golgi outposts. The multi-subunit dynein motor complex has several cofactors that are important for its activity (Reck-Peterson et al., 2018); in Drosophila neurons, this includes the conserved cofactor nudE (Arthur et al., 2015). In the absence of nudE, axons are infiltrated by Golgi outposts (Figure 2, A and B)(Arthur et al., 2015). In the axons of these *nudE*^*39A/Df*^ mutant neurons, EB1::GFP comets travel both anterograde and retrograde, which indicates a disruption in axonal microtubule polarity (Figure 2, C and D). Although EB1::GFP can also bind to slowly growing microtubule minus-ends (Feng et al., 2019), the speed of the retrograde comets in the *nudE*^*39A/Df*^ mutant axons is indicative of microtubule plus-end growth (Figure 2E). Thus, the ectopic axonal Golgi outposts in the *nudE*^*39A/Df*^ mutant neurons correlate with a change in axonal microtubule polarity.

**Fig. 2.**
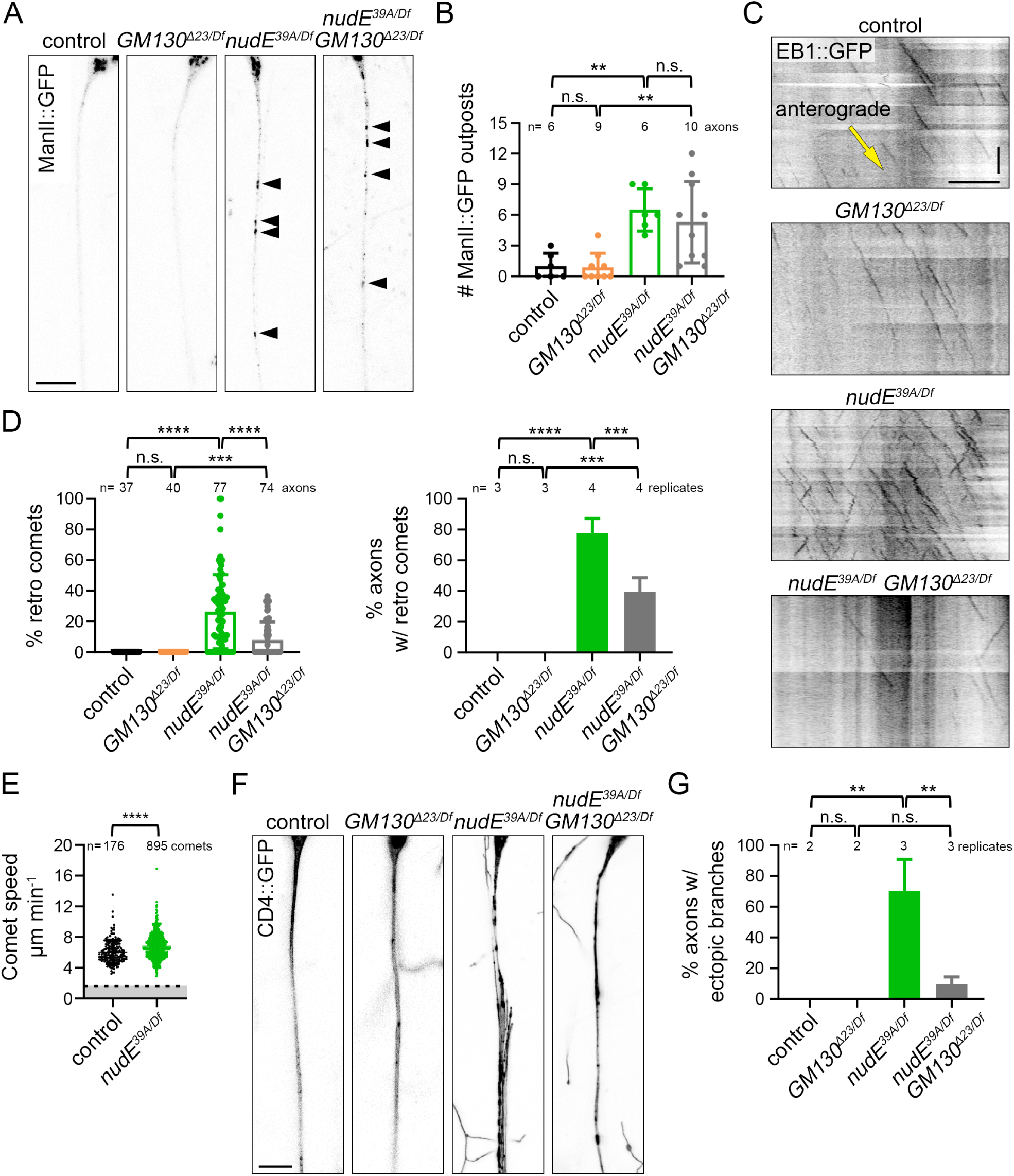
Misoriented microtubules in *nudE* mutant axons are significantly reduced when GM130 is eliminated. (A,B) Golgi outposts (marked by ManII::GFP, arrowheads) mislocalize to axons in *nudE*^*39A/Df*^ mutant neurons. The loss of GM130 does not affect the distribution of Golgi outposts in either control or *nudE*^*39A/Df*^ mutant neurons. n.s.=not significant and \*\**P*=0.01–0.001, one-way ANOVA with Tukey’s post-hoc analysis. Scale bar: 10 µm. (C,D) In the absence of nudE, axonal microtubule polarity is perturbed. Eliminating GM130 reduces the number of misoriented microtubules in *nudE*^*39A/Df*^ mutant axons. Cell body is to the left; yellow arrow indicates the direction of anterograde comet movement. n.s.=not significant, \*\*\**P*=0.001–0.0001 and \*\*\*\**P*<0.0001, Kruskal-Wallis with post-hoc Dunn’s multiple comparison analysis (% retro comets) and one-way ANOVA with Tukey’s post-hoc analysis (% axons). Scale bars: 10 µm (x-axis) and 30 sec (y-axis). Antero = anterograde, retro = retrograde. (E) The speed of EB1::GFP comets in control and *nudE*^*39A/Df*^ mutant axons is consistent with microtubule plus-end growth, indicating *nudE*^*39A/Df*^ mutant axons indeed contain microtubules with mixed polarity (comets moving at speeds below the dotted line would be consistent with microtubule minus-end growth). \*\*\*\**P*<0.0001, Student’s unpaired t test. (F,G) The ectopic branches that sprout from *nudE*^*39A/Df*^ mutant axons are suppressed by removing GM130. n=16 (control), 27 (*GM130*^*Δ23/Df*^), 59 (*nudE*^*39A/Df*^), and 41 (*GM130*^*Δ23/Df*^; *nudE*^*39A/Df*^) axons in replicates as indicated (G). n.s.=not significant and \*\**P*=0.01–0.001, one-way ANOVA with Tukey’s post-hoc analysis. Scale bar: 10 µm. All data are mean ± standard deviation.

To determine whether ectopic Golgi outposts might contribute to the alteration in axonal microtubule polarity in *nudE*^*39A/Df*^ mutant axons, we eliminated GM130. By itself, the loss of GM130 did not affect either the orientation of axonal microtubules or the localization of Golgi outposts (Figure 2, A-D). However, eliminating GM130 significantly reduced the number of misoriented minus-end-distal microtubules in *nudE*^*39A/Df*^ mutant axons (Figure 2, C and D). Golgi were still present in the *nudE*^*39A/Df*^ *GM130*^*Δ23/Df*^ double-mutant axons, indicating that the reduction in minus-end-distal microtubules was not due to a rescue of Golgi localization (Figure 2, A and B). Notably, the loss of GM130 did not completely suppress the appearance of misoriented microtubules. This may be due to the "microtubule gatekeeper" role that dynein is proposed to play in maintaining axonal microtubule polarity. In addition to carrying cargo, dynein also transports, or slides, microtubules (Rao and Baas, 2018). Dynein anchored in the proximal axon translocates microtubules into or out of the axon and prevents the entry of minus-end-distal microtubules (del Castillo et al., 2015; Rao and Baas, 2018; Rao et al., 2017). Our data suggest that dynein also maintains axonal microtubule polarity by excluding Golgi outposts that have the capacity to induce changes in microtubule organization.

The presence of ectopic Golgi outposts in the *nudE*^*39A/Df*^ mutant axons also correlates with the formation of ectopic axonal branches (Arthur et al., 2015). The axons of neurons lacking nudE develop multiple fine branches that run parallel to the main axon but terminate before reaching the ventral nerve cord; occasionally ectopic branches even extend back toward the cell body and dendrites (Figure 2, F and G). Since dendritic Golgi outposts have been correlated with dendrite branch formation and stability and loss of GM130 decreases branch number (Liu et al., 2017; Ori-McKenney et al., 2012; Yalgin et al., 2015; Ye et al., 2007; Zhou et al., 2014), we tested whether Golgi outposts might be implicated in the formation of ectopic branches that sprout from the *nudE*^*39A/Df*^ mutant axons. We found that loss of GM130 suppressed the axonal morphology defects of the *nudE*^*39A/Df*^ mutant axons, implicating the mislocalized Golgi outposts in the growth of ectopic axonal branches (Figure 2, F and G). Together, these data suggest that the ectopic Golgi outposts may be a key contributing factor to both the cytoskeletal and morphological defects of the *nudE*^*39A/Df*^ mutant axons. Thus, Golgi outposts are likely capable of inducing changes both in the microtubule cytoskeleton and neurite branching.

### Golgi outposts affect microtubule polarity independently of γ-TuRC-mediated microtubule nucleation

Dendritic Golgi outposts have been reported to serve as platforms for oriented microtubule growth during dendrite branch extension (Ori-McKenney et al., 2012; Yalgin et al., 2015). This suggests that Golgi outposts might influence microtubule polarity by controlling microtubule nucleation. Therefore, we tested whether the misoriented microtubules in the *nudE*^*39A/Df*^ mutant axons resulted from ectopic nucleation at Golgi outposts. Microtubule nucleation at Golgi membranes is templated by γ-tubulin whose nucleation activity is regulated by additional proteins, including cnn and γ-TuRC components (Martin and Akhmanova, 2018; Sanders and Kaverina, 2015; Tann and Moore, 2019). In dendrites, cnn has been implicated in regulating the directional growth of Golgi-derived microtubules during branching (Yalgin et al., 2015). Thus, we focused on whether γ-TuRC-mediated microtubule nucleation might mediate the Golgi-induced change in microtubule polarity in *nudE*^*39A/Df*^ mutant axons.

We have previously shown that reducing γ-tubulin does not suppress the appearance of minus-end-distal microtubules in *nudE*^*39A/Df*^ mutant axons (Arthur et al., 2015). Nonetheless, we followed-up our earlier findings by testing the γ-tubulin regulator cnn and the γ-TuRC component GCP4, known as dGrip75 in Drosophila (Verollet et al., 2006). Consistent with our prior report, we found that eliminating either cnn (*cnn*^*HK21/Df*^) or dGrip75 (*dGrip75*^*175/Df*^) does not suppress the formation of misoriented microtubules in *nudE*^*39A/Df*^ mutant axons (Figure 3, A and B). Moreover, we found the overall polarity of the dendritic microtubule cytoskeleton was not affected by the loss of either dGrip75 or cnn (Figure 3C). This suggests that the ectopic Golgi outposts affect microtubule polarity through a γ-TuRC-independent pathway. Nevertheless, it is still possible that Golgi outposts serve as platforms for microtubule nucleation as previously described; however, our findings suggest that the effects that outposts have on dendritic microtubule polarity would likely be limited and potentially tailored to regulate dendrite branch growth (Ori-McKenney et al., 2012; Yalgin et al., 2015; Zhou et al., 2014). Altogether, our results suggest that Golgi outposts likely alter microtubule polarity through a nucleation-independent mechanism.

**Fig. 3.**
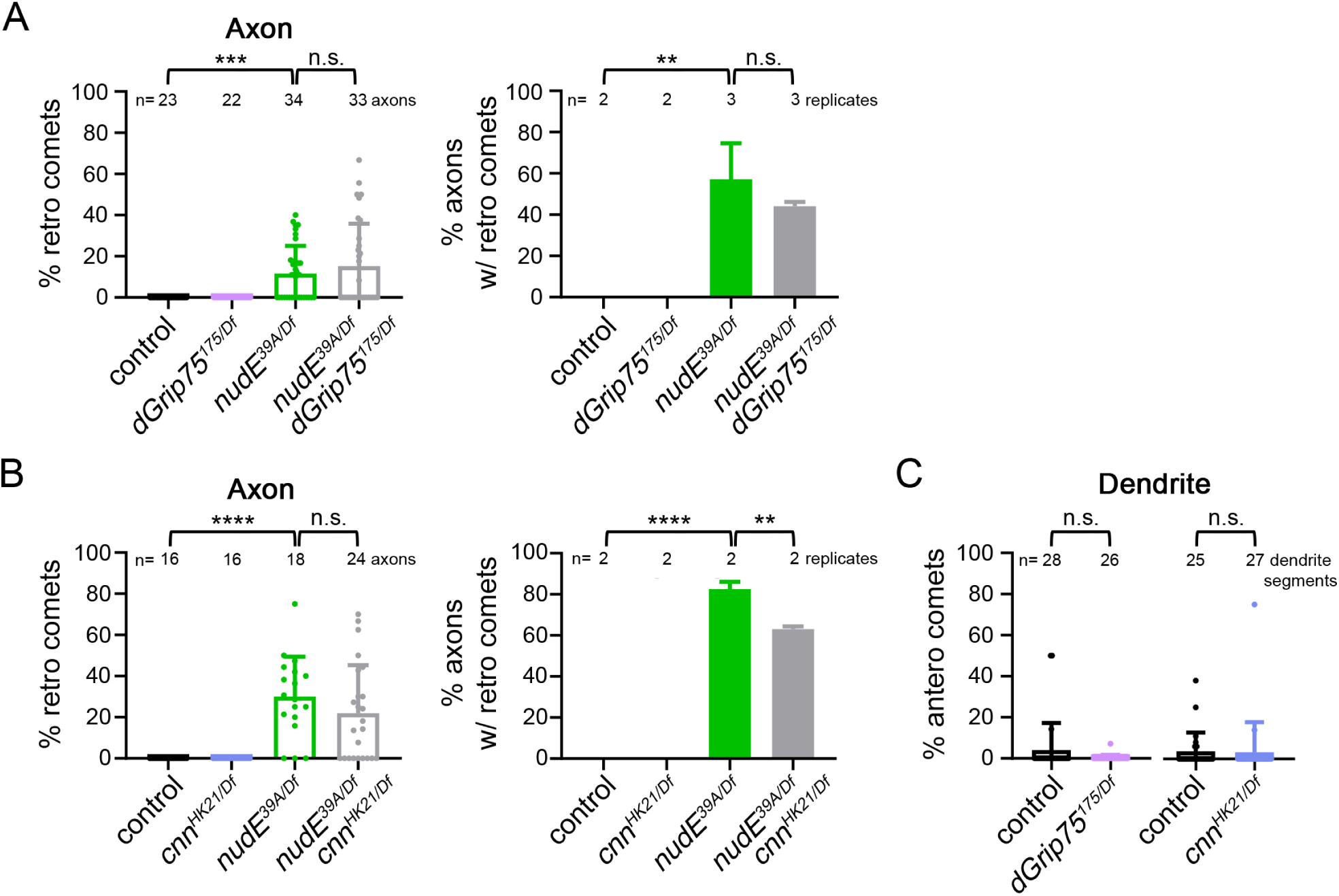
Appearance of misoriented microtubules in *nudE* mutant axons does not depend on microtubule nucleation machinery. (A,B) Eliminating either dGrip75 (A) or cnn (B) does not affect the microtubule polarity phenotype of *nudE*^*39A/Df*^ mutant axons. n=23 (control), 22 (*dGrip75*^*175/Df*^), 34 (*nudE*^*39A/Df*^), and 33 (*dGrip75*^*175/Df*^; *nudE*^*39A/Df*^) axons (A) and n=16 (control), 16 (*cnn^HK21/Df^*), 18 (*nudE*^*39A/Df*^), and 24 (*cnn*^*HK21/Df*^; *nudE*^*39A/Df*^) axons (B) in replicates as indicated. n.s.= not significant, \*\**P*=0.01–0.001, and \*\*\**P*=0.001–0.0001, Kruskal-Wallis with post-hoc Dunn’s multiple comparison analysis. (C) Microtubule polarity is normal in *dGrip75*^*175/Df*^ or *cnn*^*HK21/Df*^ mutant dendrites. n.s.=not significant, Mann-Whitney test. All data are mean ± standard deviation.

### Elevating GM130 levels increases the number of multi-compartment Golgi and alters microtubule polarity

The ability of Golgi to regulate microtubule polarity may depend on the connectedness of Golgi compartments and the formation of multi-compartment units. In addition to its role in the MTOC activity of Golgi, GM130 promotes the formation of multi-compartment Golgi units in da neurons (Martin and Akhmanova, 2018; Zhou et al., 2014). Such connectedness between Golgi compartments is thought to be necessary for the coordinated regulation of microtubules by protein complexes that are present on the *cis* and *trans* Golgi compartments (Rios, 2014; Sanders and Kaverina, 2015). We asked whether increasing GM130 levels would be sufficient to increase the number of multi-compartment Golgi outposts and to alter microtubule polarity. We found that GM130 over-expression increased the number of multi-compartment outposts in dendrites as previously reported (Figure 4A) (Zhou et al., 2014). Thus, our results provide additional support to the idea that GM130 is integral to the connectedness of Golgi compartments in neurons (Liu et al., 2017; Zhou et al., 2014); this is significant given that the role of GM130 in Golgi stack formation in other cell types has been debated (Baschieri et al., 2014; Kondylis and Rabouille, 2003; Marra et al., 2007; Puthenveedu et al., 2006; Tormanen et al., 2019). Notably, this increase in multi-compartment units correlated with an increase in anterograde EB1::GFP comets (Figure 4, A and B). This indicates that the formation of ectopic multi-compartment outposts is sufficient to alter microtubule polarity in dendrites.

**Fig. 4.**
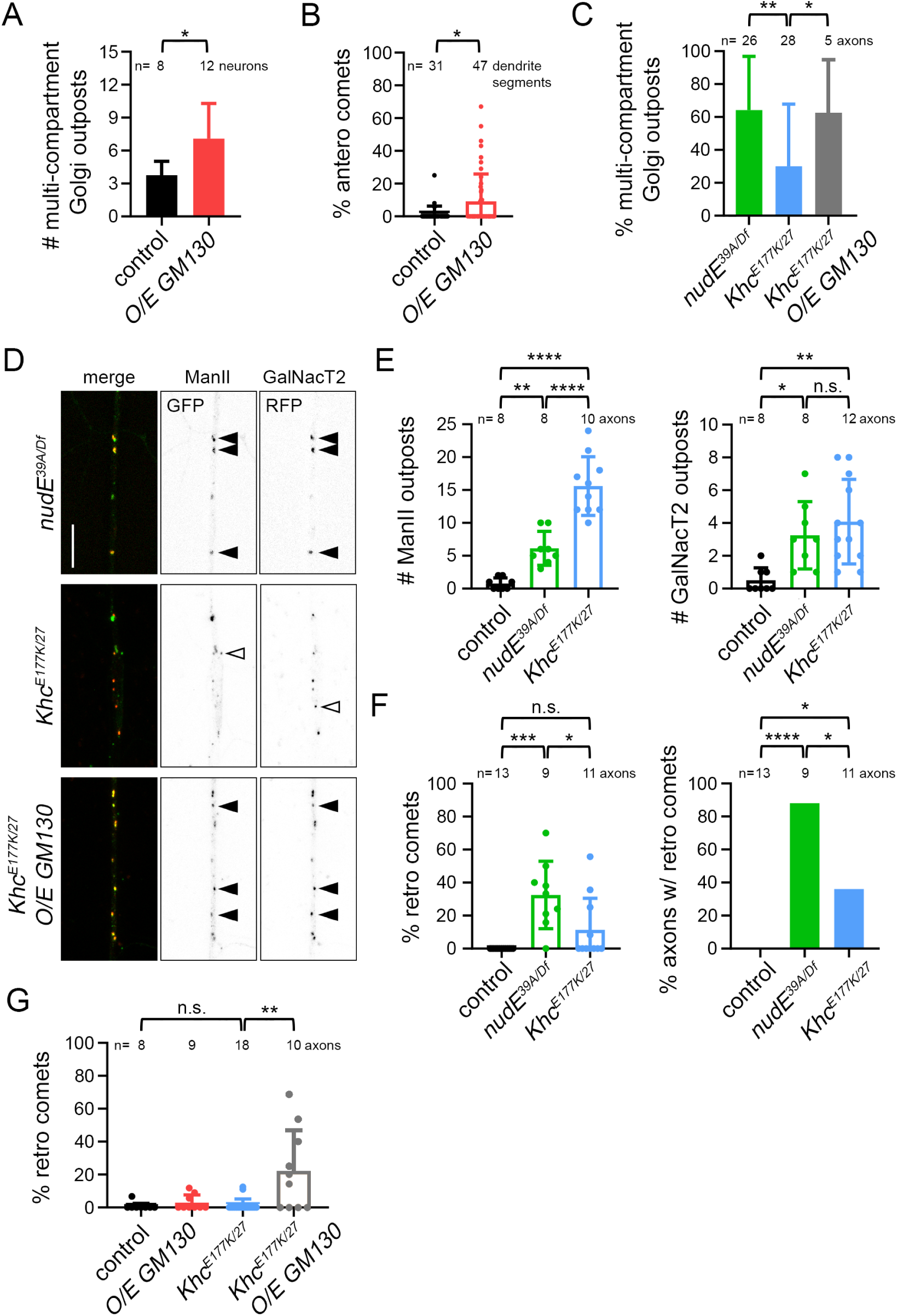
Multi-compartment Golgi are sufficient to affect microtubule polarity. (A,B) Increasing GM130 levels increases the number of multi-compartment outposts and disrupts dendritic microtubule polarity. \**P*=0.05–0.01, Mann-Whitney test. (C-E) Axons of *Khc*^*E177K/27*^ mutant neurons contain fewer multi-compartment Golgi outposts than *nudE*^*39A/Df*^ mutants, despite have similar or more numbers of ManII-and GalNacT2-positive compartments. Over-expressing GM130 increases the percentage of multi-compartment outposts in *Khc*^*E177K/27*^ mutant axons. n.s.=not significant, \**P*=0.05–0.01, \*\**P*=0.01–0.001, \*\*\**P*=0.001–0.0001 and \*\*\*\**P*<0.0001; one-way ANOVA with Tukey’s post-hoc analysis. Scale bar: 10 µm. Closed arrowheads indicate multi-compartment outposts and open arrowheads indicate single compartments. (F) In contrast to *nudE*^*39A/Df*^ mutants, *Khc*^*E177K/27*^ mutant axons have normal microtubule polarity. n.s.=not significant, \**P*=0.05–0.01, \*\*\**P*=0.001–0.0001 and \*\*\*\**P*<0.0001; Kruskal-Wallis with post-hoc Dunn’s multiple comparison analysis. (G) The increase in multi-compartment Golgi outposts in *Khc*^*E177K/27*^ mutant axons that results from the over-expression of GM130 is accompanied by an increase in misoriented axonal microtubules. n.s.=not significant and \*\**P*=0.01–0.001, Kruskal-Wallis with post-hoc Dunn’s multiple comparison analysis. All data are mean ± standard deviation.

To further test the idea that Golgi compartmentalization correlates with an effect on microtubule polarity, we turned to a mutation in *Kinesin heavy chain* (*Khc*^*E177K*^) that enhances kinesin-1 activity by disrupting motor autoinhibition (Kelliher et al., 2018). Golgi outposts mislocalize to *Khc*^*E177K*^ mutant axons; however, unlike in *nudE*^*39A/Df*^ mutant axons, the polarity of axonal microtubules is not significantly affected (Kelliher et al., 2018). Our results indicate that the compartmental organization of Golgi outposts is key to their ability to affect microtubule polarity in dendrites. This led us to characterize the compartmental organization of the ectopic Golgi outposts in the *Khc*^*E177K*^ and *nudE*^*39A/Df*^ mutant axons to determine whether the differences in axonal microtubule polarity in the two mutants might correlate with differences in Golgi compartmentalization. More specifically, we reasoned that there may be a higher number of multi-compartment Golgi outposts in *nudE*^*39A/Df*^ mutant axons, which have altered axonal microtubule polarity, than in the *Khc*^*E177K*^ mutant axons, which do not.

We analyzed the distribution of ManII::GFP and GalNacT2::TagRFP in *Khc*^*E177K*^ and *nudE*^*39A/Df*^ mutant axons. Consistent with the idea that compartment connectedness enables Golgi to influence microtubule polarity, we found that there was a higher percentage of multi-compartment Golgi outposts in the *nudE*^*39A/Df*^ mutant axons than the *Khc*^*E177K/27*^ mutant axons (Figure 4, C and D). Notably, *Khc*^*E177K/27*^ mutant axons contained equal numbers of GalNacT2-positive outposts and more ManII-positive outposts than *nudE*^*39A/Df*^ mutant axons (Figure 4E). This indicates that the *Khc*^*E177K/27*^ mutant axons have just as many Golgi compartments as the *nudE*^*39A/Df*^ mutant axons, but that the compartments are not as connected. Correspondingly, microtubule polarity is largely normal in *Khc*^*E177K/27*^ mutant axons, which is in contrast to the *nudE*^*39A/Df*^ mutant axons (Figure 4F). Combined, these results suggest that nudE (and dynein activity) are needed for the proper localization, but not the formation, of multi-compartment Golgi outposts. In contrast, enhancing kinesin-1 activity both perturbs Golgi localization and antagonizes the connectedness of Golgi compartments.

We then asked whether increasing GM130 might alter the polarity of axonal microtubules in the *Khc*^*E177K/27*^ mutant axons, which contain predominantly single-compartment Golgi outposts. In control neurons, the over-expression of GM130 alone did not affect axonal microtubule polarity (Figure 4G). The over-expression of GM130 in *Khc*^*E177K/27*^ mutant neurons both increased the percentage of multi-compartment outposts and resulted in the appearance of ectopic minus-end-distal microtubules (Figure 4, C and G). Altogether, our data support the idea that multi-compartment Golgi outposts have the capacity to remodel microtubule polarity locally even if they are not essential to the overall polarity of the dendritic cytoskeleton.

## DISCUSSION

Microtubule networks in axons and dendrites have distinct polarities. In a variety of cell types microtubule orientation is regulated by MTOCs, raising the question of whether neurons have MTOCs that carry out a similar function. In dendrites, Golgi outposts are likely candidates (Ori-McKenney et al., 2012; Yalgin et al., 2015; Zhou et al., 2014). The compartmental organization of Golgi outposts and their correlation with microtubule growth initiation sites were recently shown to depend on the *cis* Golgi matrix protein GM130 (Zhou et al., 2014). We found that eliminating GM130 does not affect the predominantly minus-end-distal orientation of microtubules in class IV da neuron dendrites, suggesting that Golgi outposts are not necessary for the unique polarity of the dendritic microtubule network. This raised the question of whether Golgi outposts might have any capacity to affect microtubule orientation. Our analysis suggests that ectopic multi-compartment Golgi outposts are sufficient to alter microtubule polarity and likely do so independently of microtubule nucleation. We propose that the unique polarity of the dendritic microtubule cytoskeleton is established and maintained independently of Golgi outposts, but that multi-compartment Golgi outposts have the capacity to locally influence microtubule polarity during events such as dendrite branch extension.

Studies carried out in mammalian cells have shown that Golgi serve as platforms for microtubule nucleation, anchoring, and stabilization (Fu et al., 2019; Martin and Akhmanova, 2018; Sanders and Kaverina, 2015). These activities arise from distinct protein complexes, whose recruitment to the *cis* Golgi depends on GM130. Golgi are generally thought to anchor and stabilize microtubules that have been generated at Golgi membranes, although the molecular mechanism by which Golgi would selectively capture these microtubules is unclear (Martin and Akhmanova, 2018; Zhu and Kaverina, 2013). Our studies indicate that γ-TuRC-mediated microtubule nucleation is dispensable for Golgi outposts to affect microtubule polarity. Consistently, a new study indicates that γ-tubulin only rarely associates with outposts (Mukherjee et al., 2019). One possibility is that microtubules are nucleated at Golgi independently of γ-tubulin. For example, it was recently reported that the mammalian tubulin polymerization promoting protein (TPPP) nucleates microtubules at Golgi outposts independently of γ-tubulin; however, TPPP is enriched in glia cells, not neurons in the mammalian nervous system (Fu et al., 2019). Another possibility is that Golgi outposts may be able to capture and stabilize microtubules that are growing in the vicinity of outposts in the relatively confined spaces of dendrites (and axons). For example, in the *nudE* mutant axons, ectopic multi-compartment Golgi outposts may perturb microtubule polarity by stabilizing nearby misoriented microtubules that would otherwise be eliminated. Thus, outposts may influence microtubule polarity by tethering and stabilizing microtubules that are not generated at Golgi.

A key component of the complexes that tether and stabilize microtubules on Golgi in vertebrate cells is myomegalin (Roubin et al., 2013; Wang et al., 2014; Wu et al., 2016). However, there is no clear Drosophila ortholog of myomegalin, making it difficult to directly test this model. The most closely related family member in flies is cnn, which is implicated in activating γ-tubulin-templated microtubule assembly (Choi et al., 2010; Roubin et al., 2013) and which we have shown is not necessary for Golgi outposts to alter axonal microtubule polarity. Moreover, new findings call into question whether cnn strongly associates with Golgi in fly neurons (Mukherjee et al., 2019). Another component of the Golgi-associated complex that anchors and stabilizes microtubules is the microtubule minus-end-binding protein CAMSAP2, whose fly ortholog is Patronin. Recent studies using da neurons have shown that Patronin is needed for minus-end-distal microtubules in dendrites, and likely acts by antagonizing the kinesin-13 microtubule depolymerase KLP10A (Feng et al., 2019; Wang et al., 2019). Our preliminary analysis of Patronin localization, however, makes it unclear whether Patronin localizes to or functions at Golgi outposts. In work using mitotic mammalian cells, GM130 has also been implicated in the stabilization of microtubules on Golgi through a mechanism that depends on the microtubule-associated protein TPX2; however, TPX2 structure and function are likely not conserved between mammals and flies (Goshima, 2011; Hayward et al., 2014; Wei et al., 2015). Thus, additional studies are needed to identify the molecular players that tether and stabilize microtubules on Golgi in Drosophila neurons.

The organization of the Golgi apparatus into a multi-compartment stack gives it a morphological and functional polarity. Correspondingly, microtubules associated with Golgi are proposed to be oriented in a particular direction relative to the Golgi compartments (Efimov et al., 2007; Martin and Akhmanova, 2018; Zhu and Kaverina, 2013). In da neurons, the correlation between microtubule polarity and Golgi compartment organization is supported by findings that microtubule growth typically initiates in a single direction from an outpost (Ori-McKenney et al., 2012; Yalgin et al., 2015). This suggests a relationship between the polarity of the Golgi stack and the associated microtubules. Thus, the relative orientation of the Golgi outpost stack likely influences the polarity of Golgi-associated microtubules (Delandre et al., 2016). Our finding that elevated GM130 levels altered microtubule polarity in dendrites may suggest that GM130 instigated the formation of misoriented Golgi stacks that in turn stabilized misoriented microtubules. Given the potential of multi-compartment outposts to influence microtubule polarity, it will be interesting to determine how Golgi outpost compartmentalization in dendrites is controlled to ensure proper dendritic microtubule polarity.

## MATERIALS AND METHODS

### Fly stocks

The following alleles and transgenic fly strains from the Bloomington Drosophila Stock Center (BDSC) and individual laboratories were used as follows: *cnn*^*HK21*^ (Megraw et al., 1999; Vaizel-Ohayon and Schejter, 1999) (BDSC 5039), which produces a drastically truncated ~106 amino acid-long protein that is not detectable by western blot, and *Df(2R)BSC306* (BDSC 23689) were used to eliminate cnn; *GM130*^*Δ23*^, a protein null allele (Zhou et al., 2014) (BDSC 65255), and *Df(2R)Exel7170* (BDSC 7901) were used to eliminate GM130; *UAS-GM130::eBFP* (Zhou et al., 2014) (BDSC 65254) and *ppk-Gal4* (BDSC 32079) were used to over-express GM130; *dGrip75*^*175*^ (Schnorrer et al., 2002) (Conduit lab, University of Cambridge; Raff lab, University of Oxford) and *Df(2L)Exel7048* (BDSC 7999) were used to eliminate dGrip75; *Khc*^*E177K*^ (Kelliher et al., 2018) was used in trans to the null allele *Khc^27^* (Brendza et al., 1999) (Saxton lab, UC-Santa Cruz); *nudE*^*39A*^ (Wainman et al., 2009) (Goldberg lab, Cornell University), a protein null allele, and *Df(3L)BSC673* (BDSC 26525) were used eliminate nudE; *plp*^*5*^ (BDSC 9567)(Martinez-Campos et al., 2004), an EMS-induced loss-of-function allele that strongly reduces plp levels, and *Df(3L)Brd15, pp* (BDSC 5354) were used to eliminate plp activity; *ppk*-*ManII::GFP* (Jenkins et al., 2017) and *UAS-GalNacT2::TagRFP* (Zhou et al., 2014) (Ye lab, University of Michigan) were used to label *medial* and *trans* Golgi compartments, respectively; *ppk-CD4::GFP* (BDSC 35842, BDSC 35843) and *ppk-CD4::tdTom* (Han et al., 2011) (BDSC 35844, BDSC 35845) were used to visualize neuron morphology; *ppk-EB1::GFP* (Arthur et al., 2015) was used to analyze microtubule polarity and dynamics.

### Live imaging

Fly crosses for live imaging were set up using 5-8 virgin females and 4-6 young males; larvae were collected in 12-h intervals and aged to the desired developmental stage (larvae produced in the first 24-48 h after mating were not used). Larvae of the desired genotype were washed with 1X phosphate-buffered saline (PBS), mounted in a 50:50 1X PBS:glycerol solution on a slide between two strips of vacuum grease, and immobilized by pressing on a coverslip mounted on top of the larva and vacuum grease spacers. The dorsal class IV ddaC neurons within abdominal segments 2-4 were imaged with a 40×1.3 NA oil immersion objective. All imaging was performed on a Leica SP5 (Leica Microsystems) using HyD photodetectors.

### Microtubule polarity and growth analysis

Microtubule polarity and growth were analyzed by scoring EB1::GFP comets within 150 µm of the cell body in dendrites or axons. EB1::GFP comet trajectories were captured at a resolution of 1024 x 512 pixels and a rate of 0.86s per frame for 5-7min. One or two ddaC neurons per larva were imaged at 96 −120 h after egg laying (AEL). Videos were stabilized with the stabilizer plugin in FIJI ImageJ (ImageJ; National Institutes of Health); kymographs were generated in Metamorph (Molecular Devices). Comet trajectories were manually traced, and the position and time coordinates were recorded to calculate comet direction (microtubule orientation) and frequency. A comet trajectory was included only if it could be clearly traced in at least 12 continuous frames (~11s). Anterograde comets traveled away from the cell body whereas retrograde comets traveled toward the cell body. The frequency of EB1::GFP comets was calculated as the number of comets present in a 100 µm segment (axons) per minute. To calculate microtubule polarity in dendrites, EB1::GFP comets were scored in a segment ≥ 30 µm between two branchpoints. Movies that did not contain at least two comets were excluded.

### Axon branching analysis

Axon morphology was visualized at late 3^rd^ instar (120-144h AEL) using CD4::GFP and CD4::Tomato. Images were captured at 1024×1024 resolution and 1 µm z-steps (5-15 steps total). Analysis was performed on axons within 150 µm of the cell body. Z-stacks 5-15 µm thick were max projected for analysis. Axon branching was assessed manually by determining whether an axon split into one or more branches.

### Golgi compartment analysis

Images of ManII::GFP and GalNacT2::TagRFP puncta in the dendrites and axons of neurons in 96-120h AEL larvae were captured at a resolution of 1024 x 1024 pixels and 0.75 µm z-steps over 5-15 µm. Analysis of Golgi outposts in axons included outposts within 100 µm of the cell body. For dendrite analysis, Golgi outposts throughout the entire arbors were included and split into two groups: (1) those within a radius of 95 µm of the cell body and (2) those outside this radius. One to two ddaC neurons were imaged per larva. Analysis was performed on the max-projected images. Signal outside the regions of interest (ROI) in axons and dendrites were masked. Masked images were subjected to a threshold gray value of 100 for segmentation. The segmented signals were quantified with the ImageJ particle analysis function with the size cutoff of 0.10-15 µm^2^. Puncta outlines were saved as ROIs. The resulting particle numbers and sizes were exported to Excel for analysis. For multi-compartment analysis, overlapping compartments were manually scored by overlaying the puncta outlines from each channel, namely ManII::GFP and GalNacT2::TagRFP. Golgi units were scored as multi-compartment when the ManII::GFP and GalNacT2::TagRFP signal overlapped.

### Statistical analysis

Statistical analyses were performed using GraphPad Prism8. The Anderson-Darling and Shapiro-Wilk tests were used to determine whether data were normally distributed. For normally distributed data, Student’s unpaired t tests were used to compare two groups; one-way ANOVA with post hoc Tukey was used for multiple comparisons. For non-normally distributed data, Mann-Whitney’s test was used to compare two groups and Kruskal-Wallis test followed by post hoc Dunn’s was used for multiple comparisons. Fisher’s Exact test was used for comparing proportions. P=0.05 was used as a cutoff for significance. Significance levels are represented as: \**P*=0.05–0.01, \*\**P*=0.01–0.001, \*\*\**P*=0.001–0.0001 and \*\*\*\**P*<0.0001. n.s.=not significant.

## Acknowledgments

We thank the following laboratories and stock centers for generously sharing Drosophila strains: Drs. Paul Conduit (University of Cambridge) and Jordan Raff (University of Oxford) for *dGrip75* flies, Bing Ye (University of Michigan) for *UAS-GalNacT2::TagRFP* flies, and the Bloomington Drosophila Stock Center (supported by NIH P40OD018537). We thank Erik Dent and Mary Halloran (University of Wisconsin-Madison), Adrian Moore (Riken CBS), Paul Conduit (University of Cambridge), and members of the Wildonger lab for insightful discussions and feedback; and Lindsay Mosher and Dena Johnson-Schlitz for technical assistance. This work is supported by the National Institutes of Health grant R01NS102385.

## Abbreviations used

cnn: centrosomin
da: dendritic arborization
EB1: End-binding 1
γ-TuRC: γ-tubulin ring complex
Khc: kinesin heavy chain
MTOC: microtubule-organizing center
plp: pericentrin-like protein

